# The Good, the Bad, and the Ugly: Segmentation-Based Quality Control of Structural Magnetic Resonance Images

**DOI:** 10.1101/2025.02.28.640096

**Authors:** Robert Dahnke, Polona Kalc, Gabriel Ziegler, Julian Grosskreutz, Christian Gaser

## Abstract

The processing and analysis of magnetic resonance images is highly dependent on the quality of the input data, and systematic differences in quality can consequently lead to loss of sensitivity or biased results. However, varying image properties due to different scanners and acquisition protocols, as well as subject-specific image interferences, such as motion artifacts, can be incorporated in the analysis. A reliable assessment of image quality is therefore essential to identify critical outliers that may bias results. Here we present a quality assessment for structural (T1-weighted) images using tissue classification. We introduce multiple useful image quality measures, standardize them into quality scales and combine them into an integrated structural image quality rating to facilitate the interpretation and fast identification of outliers with (motion) artifacts. The reliability and robustness of the measures are evaluated using synthetic and real datasets. Our study results demonstrate that the proposed measures are robust to simulated segmentation problems and variables of interest such as cortical atrophy, age, sex, brain size and severe disease-related changes, and might facilitate the separation of motion artifacts based on within-protocol deviations. The quality control framework presents a simple but powerful tool for the use in research and clinical settings.

## Introduction

Multicentre *magnetic resonance imaging* (MRI) studies and data-sharing projects have become increasingly common in cognitive and clinical neuroscience in recent years. The collaboration of several imaging centers, allowing for increased statistical power through larger sample sizes, is especially beneficial for investigating rare diseases and individual differences (Bethlehem et al., 2022; Markiewicz et al., 2021; Wiseman et al., 2019). However, project deviations from the initial research plans (e.g., switching from functional to a structural imaging focus), differences and changes of imaging hardware and software, quality assurance procedures, and the resulting image quality variations may introduce bias in subsequent image processing and statistical analysis (Ai et al., 2021; Bottani et al., 2022; Kruggel et al., 2010; Reuter et al., 2015). In particular, the presence of noise, (motion) artifacts, inhomogeneity, or reduced resolution could affect image processing, even when such interferences are modeled and partially corrected during data processing (AubertBroche et al., 2006; Nárai et al., 2022; Tian et al., 2021; Figure 1).

**Figure 1:**
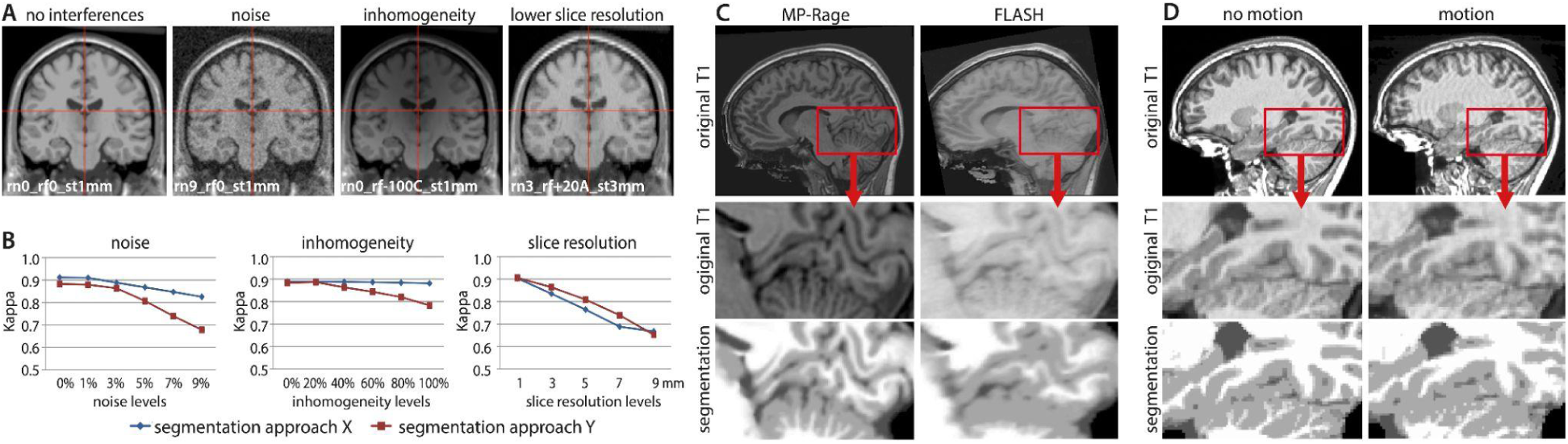
(A) Image properties such as noise, inhomogeneity, and resolution influence the segmentation accuracy. (B) The segmentation accuracy can be quantified by the kappa similarity statistic (Cohen, 1960), here presented for two segmentation approaches on simulated images (AubertBroche et al., 2006). (C) Shows an illustration in real data with reduced anatomical details in a FLASH protocol (Kempton et al., 2011) or (D) in case of movement artifacts (MR-ART sub-988484 from Nárai et al., 2022).

Typically, manual *quality control* (**QC**) checks each image for scan-specific interferences (e.g., motion artifacts) by visual inspection to remove outliers (Keshavan et al., 2018, Kruggel et al., 2010; Nakua et al., 2023). However, manual assessment is time-consuming, highly subjective, and typically relies on project-specific definitions (Nárai et al., 2022). To make this process more efficient and reliable, automated quality control approaches have been proposed for structural (Bottani et al., 2022; Esteban et al., 2017; Mortamet et al., 2009), functional (e.g., Christodoulou et al., 2013), and diffusion imaging (e.g., Maximov et al., 2021). In addition, the image quality estimates can be used to harmonize imaging data (Garcia-Dias et al., 2020; Lutti et al., 2022; Pomponio et al., 2019). A systematic overview of different QC frameworks has been provided by Hendriks et al. (2024).

In this study, we propose a powerful and easily applicable QC framework for structural (T1-weighted) MRI data. Earlier versions have been extensively evaluated in Gilmore et al. (2021) and Ma et al. (2022). The proposed QC framework introduces, standardizes and integrates different quality metrics into a continuous *structural image quality rating* (**SIQR**). It supports both automatic and interactive assessments of a preprocessed MRI scan’s suitability for prospective use, as well as the identification of potential outliers within a sample, ensuring unbiased data analysis. All measures and tools are part of the *Computational Anatomy Toolbox* (**CAT;** https://neuro-jena.github.io//cat, Gaser et al., 2024) of the *Statistical Parametric Mapping* (**SPM;** http://www.fil.ion.ucl.ac.uk/spm, Ashburner et al. 2002) software and also available as a standalone version (https://neuro-jena.github.io/enigma-cat12/#standalone).

## Methods

This section presents the rationale for a segmentation-based QC framework, a definition of several quality measures, their standardization into quality scales and the integrated composite measure *Structural Image Quality Rating* (**SIQR**). We further describe the detection of imaging artifacts based on the within-sample quality and introduce the interactive outlier detection. Finally, we evaluate the proposed measures using simulated and real MRI data.

### Segmentation-based Image Quality Assessment

For practical reasons, our QC framework uses the raw NIFTI format rather than the original DICOM format, as NIFTI images are more commonly available in public datasets and are more often used as input in data processing tools (Markiewicz et al., 2021). The QC framework relies on an existing conventional or deep-learning based classification of brain tissues, which is usually a prerequisite for subsequent brain image analyses (e.g., Ashburner et al., 2005, Gaser et al., 2024, Mendrik et al., 2015, Billot et al., 2023). All proposed measures are based on image properties primarily within the brain because the background might be affected by anonymization, noise or artifacts (Figure 2A; Kruggel et al., 2010; Marques et al., 2010). The quality measures are optimized to avoid the evaluation within parts of the brain that are typically affected by aging-related tissue changes, such as white matter hyperintensities, small vessel disease and perivascular spaces (Figure 2B). Within the CAT12 toolbox, the QC is the final step of the preprocessing and extends the processing time of a subject by only a few seconds. Alternatively, it can be run separately as a SPM batch for a pre-existing tissue segmentation provided by other algorithms.

**Figure 2:**
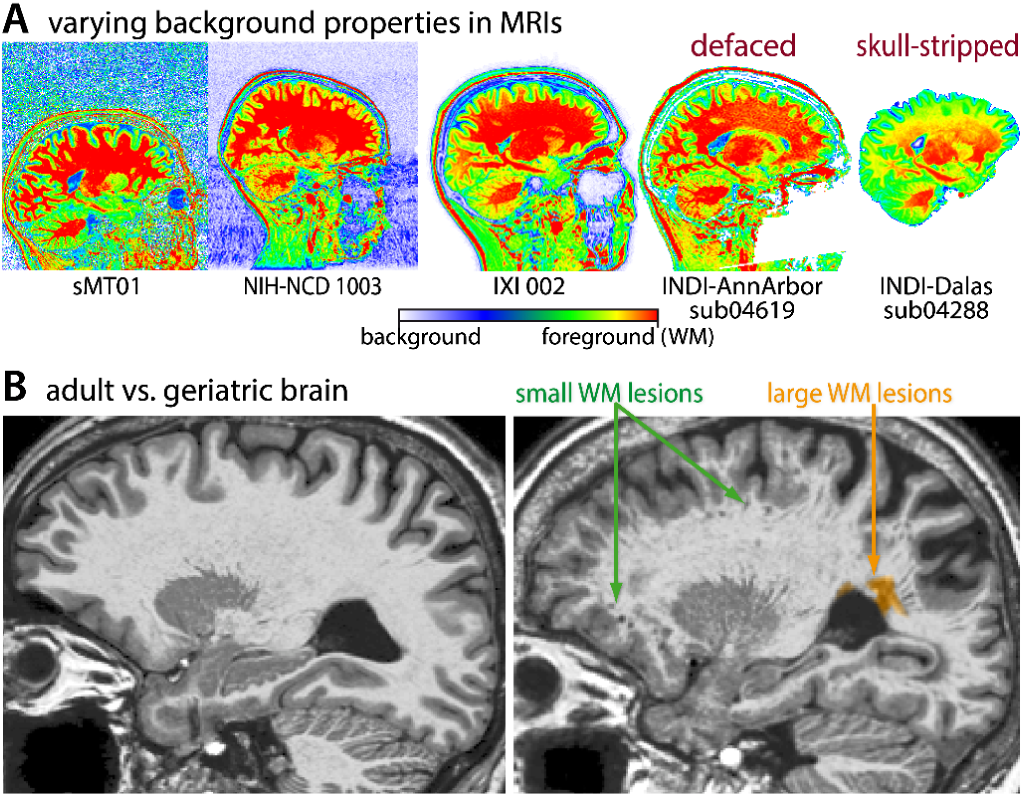
(A) Due to several different background types in real samples, only the brain tissues (excluding the background) were used for the evaluation of image quality. (B) To avoid side effects from age-related changes in volume and structure, the tissue segmentation is optimized to avoid tissue boundaries and perivascular spaces by morphological operations and masking.

The primary evaluation of our proposed QC measures is based on the *Brain Web Phantom* (**BWP**, AubertBroche et al., 2006), a simulated MRI dataset, which presents a well-established standard in developing and comparing processing methods for typical image properties such as noise, inhomogeneities, and resolution (Figure 1B).

### Quality Measures

As our measures have been optimized for use in cognitive and clinical neuroscience studies, the presentation is focused on practicality. A full (technical) description can be found in the <u>Supplemental</u> material. For intensity-based measures, we use measure-to-contrast ratios instead of contrast-to-measure ratios. This approach ensures that the ratings follow a linear scale rather than a logarithmic one, as defined by the Brain Web Phantom.

Several key quality metrics are considered:

- **Noise-to-contrast ratio (NCR)**: This metric estimates image noise by calculating the lowest average local standard deviation of voxel intensities in the bias-corrected image. It is assessed within optimized cerebrospinal fluid (CSF) and white matter (WM) regions.
- **Inhomogeneity-to-contrast ratio (ICR)**: This measure evaluates intensity variations across the image by calculating the global standard deviation of smoothed intensities within the optimized WM segment.
- **Resolution score (RES)**: To account for distortions due to anisotropic resolution, this score is directly computed using the root mean square (RMS) equation.
- **Edge-to-contrast ratio (ECR)**: Since resampling or smoothing can degrade voxel resolution, we suggest an additional measure which captures the average slope of intensity changes at the gray matter (GM)/WM boundary. This helps assess the sharpness of tissue interfaces.
- **Full-brain Euler characteristic (FEC)**: This metric quantifies the topological integrity of the WM brain interface, helping to detect potential distortions caused by noise and artifacts.

These measures provide a comprehensive assessment of MRI image quality, ensuring that intensity-based distortions, resolution issues, and structural inconsistencies are identified and accounted for.

### Standardization of Measures

Standardization into a normative range can enable simpler comparison across studies and support easier interpretations. To accommodate various international rating systems, we have adopted a linear percentage and a corresponding (alpha-)numeric scaling. (Figure 3, QR_percentage_ = 105 - QR_grade_ * 10, QR_grade_ = (105 - QR_percentage_) / 10). The quality rating ranges from 0.5 (100 *rating points* (**rps**); grade A^+^) to 10.5 (0 rps; grade F) for highest and lowest image quality, respectively. Numerical values provide a specific rating, whereas letters describe quality ranges, e.g., grade A describes values between 90 and 100 rps. Scaling of the quality measures was performed using the first half of the BWP dataset (see <u>Evaluation Concept and Data</u>), while the second half was used for evaluation. Although the BWP does not include the simulation of motion artifacts, these are in general comparable to an increase of noise in the BWP dataset by 2 percentage points. In our QC measures, this roughly corresponds to an increase of +1 grade or -10 rps compared to motion-free data. For improved (human) readability, we standardized all measures by applying a simple linear scaling function

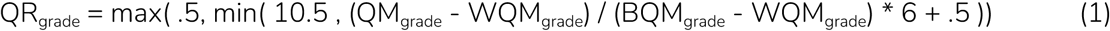

**Figure 3:**
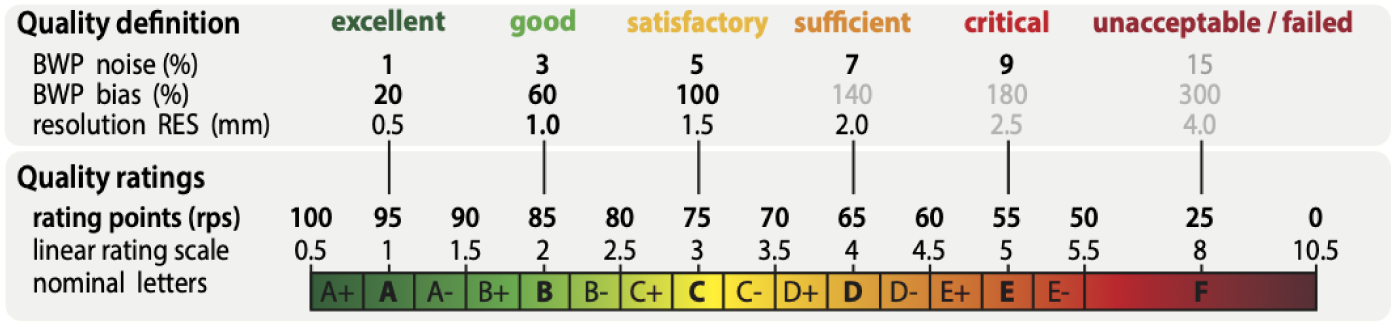
Quality rating system: The percentage, numerical, and character grades were scaled on the basis of the BWP, which represents a standard for evaluation of image processing methods. It should be noted that excellent ratings are reserved for images with exceptional quality, whereas typical scientific data generally receives “only” good assessments.

to transform the original quality measure QM into a quality rating QR, with BQM as the best (95 rps, grade 1) and WQM (45 rps, grade 5) as the worst regular value.

### Integrated Structural Image Quality Rating

The structural image quality rating (SIQR) is defined using an exponentially weighted average of multiple quality scores (see Equation 1 below). This single composite score integrates various aspects of image quality, providing a robust metric for assessing structural image quality and identifying potential outliers.

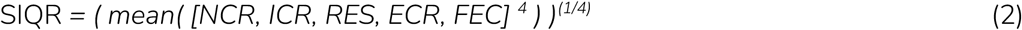

To balance the sensitivity to different quality measures while ensuring that the necessary quality conditions are met, we apply an exponentially weighted averaging approach — similar to the root mean square (RMS) but using the fourth power and fourth root. This method allows well-rated images to contribute positively without overshadowing critical quality constraints.

### Sample Normalization for Outlier Detection

So far, each image has been evaluated individually, enabling the detection of outliers with very low resolution or high noise (e.g., those falling below a C rating, such as SIQR < 70 rps). However, to identify more subtle issues—such as mild motion artifacts—it is necessary to assess deviations from the ideal quality expected for a given MRI protocol within a specific sample. To achieve this, we estimate the upper quartile of the SIQR percentage scores from images acquired with the same protocol and apply a linear correction (a simple translation, as the values have already been scaled according to the BWP). This normalization results in a standardized SIQR, where values close to zero indicate optimal protocol quality, while higher values highlight potential outliers. To establish a general threshold for detecting quality issues, we employed a *receiver operating characteristic* (**ROC**) curve analysis combined with 2-fold cross-validation (splitting the dataset into odd- and even-numbered files based on filenames). This approach was validated using test samples with expert ratings, ensuring robust performance in identifying suboptimal images.

### Software

*The “Check Sample Homogeneity*” tool (Figure 4) in the CAT12 toolbox supports a guided analysis of large datasets to detect and exclude outliers in anatomy, preprocessing and image quality from analysis by estimating a sample-specific z-score. The tool has been designed in an interactive format with the intention to encourage users to get in touch with their data and carefully decide on the in-/exclusion of the images from analyses. Artifacts can result in a systematic bias, often resulting in an underestimation of GM (Reuter et al., 2015). To ensure the validity of statistical analyses, it is suggested that severe image quality-related outliers are excluded based on normative assessments provided by the toolbox. The quality estimation is also available as *“Image Quality Estimation*” SPM batch to process selected raw (co-registered) structural scans with a given brain tissue segmentation (e.g., from SPM). The results for each input image are stored in an XML file and can be used for subsequent analysis steps and potential analysis in relation to effects of interest of a study (such as age).

**Figure 4:**
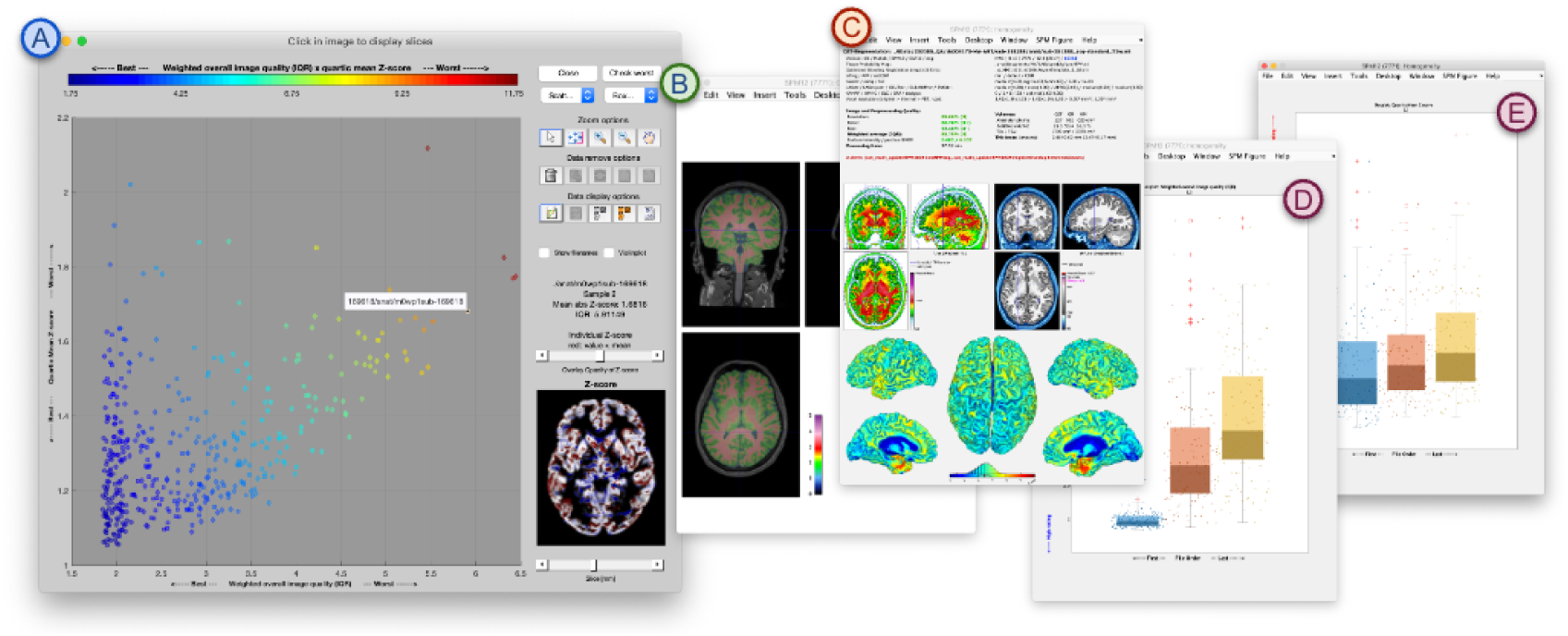
Shown is the “Check Covariance” Tool in CAT12 for the MR-ART dataset grouped by the amount of motion (see D and E). Scans can be selected in the main window (A) to understand deviating ratings by viewing the original image (with segmentation overlay; B) or the preprocessing report (C) to remove outliers with image- and processing-related problems or atypical anatomical features.

### Evaluation Concept and Data

The training and testing of our proposed measures was done using simulated images from the *Brain Web Phantom* (BWP; AubertBroche et al., 2006) and the *cortical aging phantom* (**CAP**; Rusak et al., 2021), as well as real data from IXI, ATLAS, MR-ART (Nárai et al., 2022), and a test-retest dataset (Table 1).

**Table 1:** Short overview of the used datasets.

| Dataset | *n* | Age [years] | Sex [%men] | Sites | Description |
| --- | --- | --- | --- | --- | --- |
| BWP | 600 | ~30 | 100% | 1 | Simulated dataset (AubertBroche et al., 2006; https://brainweb.bic.mni.mcgill.ca/) for basic definition (train: odd files) and evaluation (test: even files) of the quality ratings, with 5 noise levels (1% to 9%), 3 different bias fields with 5 levels (20% to 100%) and 8 resolution levels with a voxel-length of 1 and 2 mm. |
| BWPE |  | ~30 | 100% | 1 | BWP with simulated skull-stripping and segmentation errors (see Figure 5) to test the robustness of our measures in case of severe processing problems. |
| CAP | 400 | 39.1-78.3 (70.7±5.4) | 50% | 1 | Simulated atrophy dataset (Rusak et al., 2021) to test for side effects of GM tissue atrophy that correspond to aging of about 100 years. |
| IXI | 554 | 20-86 (48.5±16.4) | 44.8% | 3 | Brain aging sample to test the effects of age, brain size, and sex (only scans with complete phenotypic data; http://www.brain-development.org/). |
| ATLAS (R1.2) | 304 | NA | NA | 11 | T1 images of subjects with (masked/unmasked) lesions to test the stability in case of severe structural changes (https://fcon_1000.projects.nitrc.org/indi/retro/atlas.html). |
| MR-ART | 148*3 | 18-75 (30.0±12.8) | 35.1% | 1 | Dataset without and with intended motion artifacts (Nárai et al., 2022; https://openneuro.org/datasets/ds004173/versions/1.0.2). |
| TRT | 6 (127) | ~30 | 100% | 1 | Various MR protocols with a scan time duration from 30s to 11 minutes on a 3T Philips scanner (Thyreau et al., 2013). |

The BWP-training data consisted of all odd files (ordered by filename) and was used to scale the quality measures and estimate the weighted averaging described above. The test subset (even files) was used to quantify the relationship between quality ratings and segmentation accuracy. Moreover, we used the BWP to further simulate typical brain-extraction and segmentation artifacts (e.g., by erosion/dilation of tissue segments) to test the robustness of the quality measures in case of critical data conditions (see <u>Supplement Segmentation Issues</u>, see Figure 5). The *cortical aging phantom* described in Rusak et al. (2021) was used to test the effects of brain atrophy of up to 1 mm on our quality measures.

**Figure 5:**
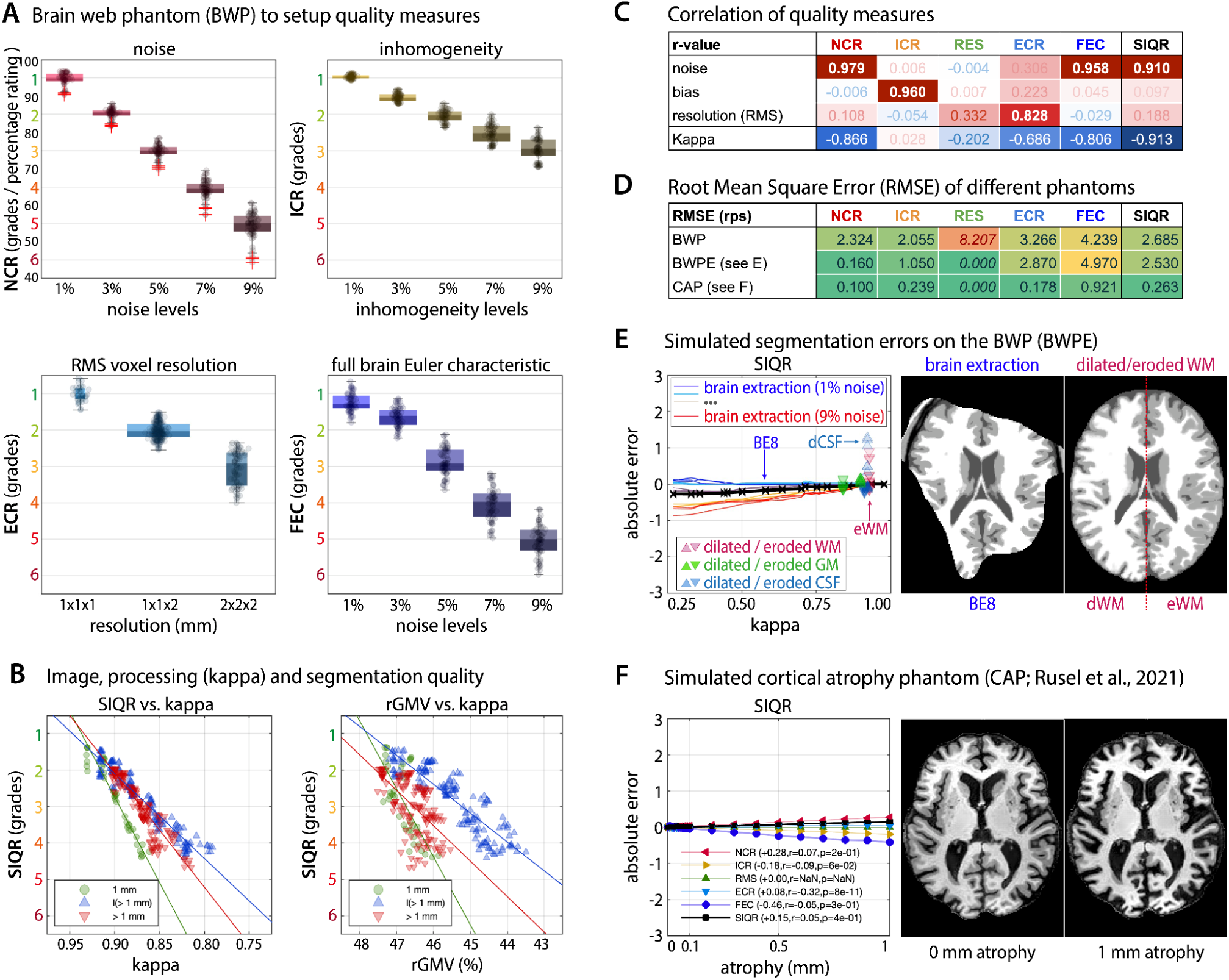
(A) Shown is the dependency of quality measures on manipulated level of noise, inhomogeneity and resolution levels using the Brain Web Phantom (BWP): NCR (noise-to-contrast rating), ICR (inhomogeneity-to-contrast rating), ECR (edge-to-contrast rating), and FEC (full brain Euler characteristic). (B) The structural image quality rating (SIQR) integrates all these measures into one score and shows significant associations with segmentation quality (characterized by kappa) and the relative gray matter volume (rGMV) based on CAT12. (C) Overall, our ratings show specific relationships to their corresponding BWP perturbations but not others and (D) small root mean square errors (RMSE) also in the case of simulated segmentation errors (E) or aging (F). Of note, the numerical grading system and percentage system are inversely scaled, where –10 rps correspond to +1 grade and roughly correspond to the emergence of obvious motion artifacts.

Although simulated data enables basic evaluation under defined conditions, real data is essential to investigate possible dependencies/biases. The measures were quantified in IXI and ATLAS datasets to test for possible effects of age, sex, and lesions. Finally, the MR-ART dataset with 148 subjects, each with 3 scans without, with light, and with severe motion artifacts, and expert ratings was used to validate the use of our measures to separate images with motion artifacts in a split-half test design.

Additionally, we used the Tohoku *test-retest* (**TRT**) dataset, which contains 126 T1-weighted scans (Thyreau et al., 2013). All scans were preprocessed, registered, and resliced to a high-resolution template with 0.50 mm isotropic resolution. A median template was used to remove outliers and to create the final ground truth segmentation by averaging. Finally, six scans were selected based on their scan time and image characteristics to evaluate the influence of image quality on segmentation accuracy.

## Results

The quality scores were first evaluated on the simulated test data in order to determine the accuracy of interference quantification and to investigate how robust the measures are in cases of simulated segmentation problems and aging. Furthermore, we used the IXI and ATLAS datasets to study the effects of aging, sex, brain size, and stroke lesion on our proposed measures of image quality. Additionally, we evaluated the ability to detect images with motion artifacts on the MR-ART dataset and demonstrate the application in a test-retest scenario. All measures had been standardized (see Figure 3) and evaluated on the BWP before focusing on the averaged SIQR score. Of note, obvious subject/scan-specific motion artifacts generally increase the scans’ rating for about 1 grade, which corresponds to a decrease of 10 rps (and +0.5 grade / -5 rps for light artifacts), in comparison to the typical rating achieved by the majority of scans of the same protocol. Similarly to the method section, we focus here on the results pertaining to SIQR and refer the interested reader to the supplementary section <u>S2</u> for a more detailed overview.

### Simulated data

The evaluation on the BWP test dataset showed that most quality ratings have a very high correlation (rho > .950, *p* < .001) with their corresponding perturbation and a very low correlation (rho < |0.1|) with the other tested perturbations (see table in Figure 5A & C). This suggests considerable specificity of the proposed quality measures. The combined SIQR score also showed a very strong association with the segmentation quality kappa (rho = –.913, *p* < .001) and brain tissue volumes (rho_CSF/GM/WM_ = –.472/–.484/.736, *p*_CSF/GM/WM_ < .001) (Figure 5B). The root mean square errors (**RMSE**) between the expected and measured values of the SIQR were 2.685, 2.530, and 0.071 rps for the BWP testset, the BWP-derived segmentation error testset, and the cortical atrophy phantom, respectively (Figure 5D).

Most notable is the quantification of the image resolution, where the simple voxel-based resolution rating RES did not work well in interpolated data (i.e., as expected in 225 out of 625 cases), resulting in a lower correlation (rho = .332) and a high RMSE of 8.207 rps. The edge-based resolution measure ECR, on the other hand, generally performed better (rho = .828, *p* < .001), but was more affected by noise (rho = .306, *p* < .001) and inhomogeneity (rho = .223, *p* < .001) than other scores. Overall, low quality data was typically found to have higher error rates than high quality images, i.e., the RMSE of the SIQR for images with <5% noise was 2.079 rps compared to

3.417 rps for images with >5% noise. The tests with simulated segmentation errors suggested that NCR and ICR were extremely robust (Figure 5E), whereas ECR and especially FEC were quite sensitive to strong (i.e., 1 voxel) over-/underestimations of CSF and WM.

### Real data

The real data analysis of IXI and ATLAS cohorts suggests that the proposed quality measures were not affected by *total intracranial volume* (**TIV,** *r*_SIQR_=.257, *p*_SIQR_=.223), sex (Mann-Whitney-U-Test: U = 38097, Z=.691, *p*=.490), or stroke lesions in the ATLAS dataset (Figure 6B), while age showed minor effects (*r*_SIQR_=.116, *p*_SIQR_=.015; Figure 6A) in IXI. Since IXI rather contains scans without significant motion artifacts, it provides a useful estimate of the typical overall variability in terms of the standard deviation of the SIQR score with 1.594 rps (Guys/HH/IOP = 1.601/1.676/1.505 rps).

**Figure 6:**
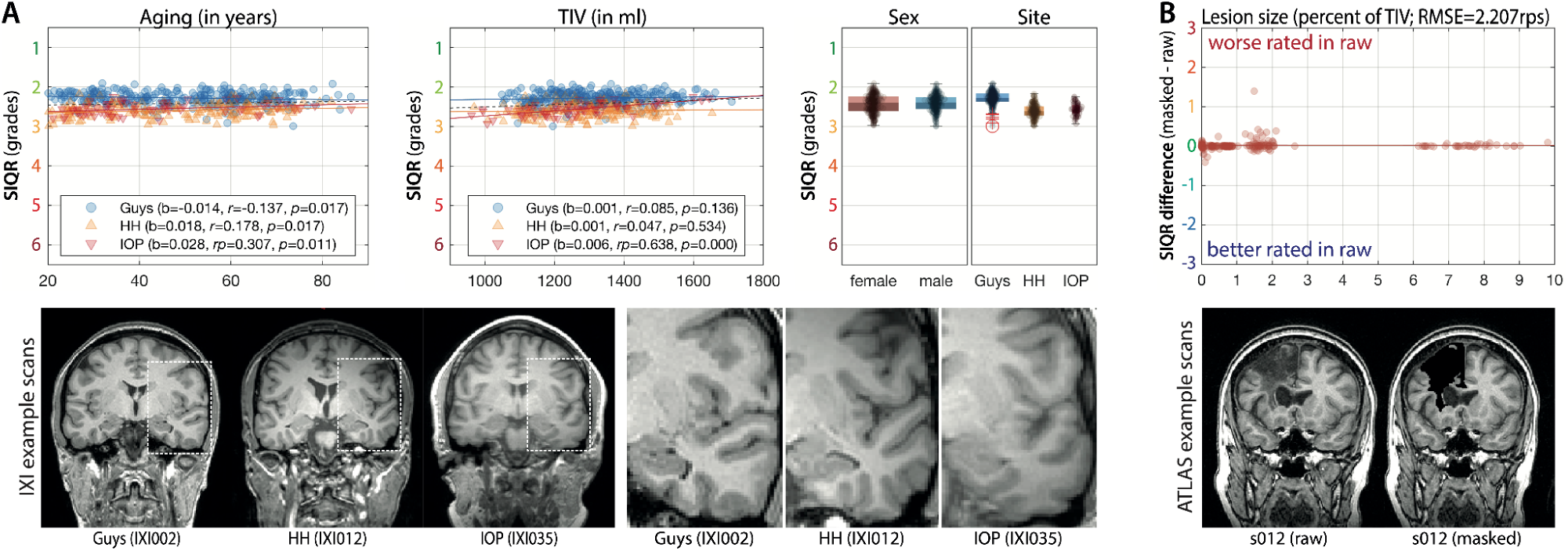
(A) The results of the structural dependency test in IXI dataset showed that the SIQR measure is independent from sex, and only slightly associated with TIV and age (see the supplement for other quality measures) with an average standard deviation of 1.594 rps per site. (B) The results from the ATLAS dataset suggest that severe structural changes in terms of lesions do not significantly affect the SIQR measure when comparing raw vs. masked images.

The effects of motion artifacts were evaluated using the MR-ART dataset (Figure 7A), which also allowed the comparison to the quality measures of MRIQC 0.16.1 (Esteban et al., 2017). In order to detect motion artifacts, each score was normalized (by subtracting the first quartile value to consider the typical protocol quality) and a *Receiver Operating Characteristic* (**ROC**) was applied. The measures were tested under 3 conditions, namely comparing (i) no vs. severe artifacts, (ii) no vs. light+severe artifacts, and (iii) no+light vs. severe artifacts (Figure 7B). The best ROC thresholds to separate good from bad scans in the 3 groups were 4.90/7.20/8.35 rps for the normalized SIQR and standard deviations of 1.441 and 3.041 rps in the no-artifact and no+light-artifact groups, respectively. The accuracy of the SIQR as determined by the ROC (an average over the three groups) was 0.889 and 0.813, with an *area under curve* (**AUC**) of 0.970 and 0.913 for CAT12 and SPM12, respectively. This was slightly better than the best performing alternative metrics from MRIQC measures especially *snr_wm*, *snr_total, cjv* and *cnr* (see Esteban et al., 2017) that achieved average accuracies over the three groups of 0.878/0.879/0.850/0.845 and an AUC of 0.961/0.952/0.937/0.930, respectively. Moreover, the 3 groups of scans of MR-ART support the estimation of the segmentation error kappa of the scans with motion compared to the motion-free scans in comparison to aging defined by SPM12 and CAT12 segmentation.

**Figure 7:**
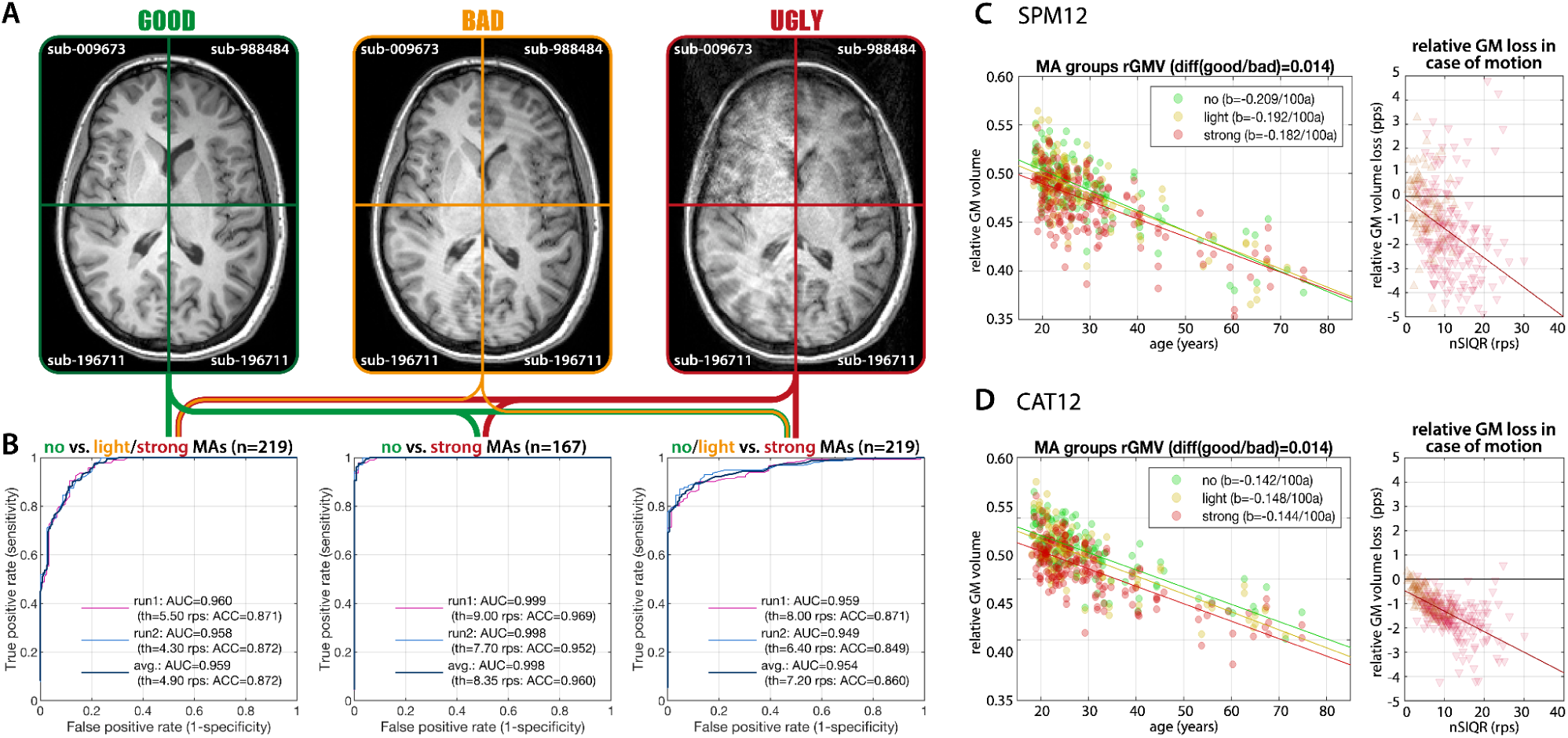
(A) Example images from the MR-ART dataset (Nárai et al., 2022) for three different conditions based on expert ratings splitting data into (no), (light), and (strong) motion artifacts. (B) Shows the Receiver Operating Characteristic (**ROC**) curves when classifying these groups using SIQR (see the Supplement for ROCs of other quality ratings) with high accuracy (**ACC**) and area under curve (**AUC**). (C & D) We further evaluated the relative GM tissue volume change in percentage points (pps) of scans with motion artifacts compared to the ones without for each subject processed by SPM12 and CAT12. The results suggest that the expected lower segmentation accuracy for lower image quality might lead to GM under- and WM overestimation.

Finally, we validated the proposed quality metrics using a scan-rescan test including six images consisting of a series with gradually increased quality (and scan-time) as well as the ground truth image (Figure 8). The expected improvement of image quality was clearly observable in terms of sharper anatomical details and reduced noise. The kappa indices, rGMVs, and SIQR scores confirmed these visual observations, but pointed out further interesting details. The noisy short-time scans S1 and S2 showed significantly lower kappa, worse SIQR ratings, and smaller GM volumes, whereas the other scans were highly comparable.

**Figure 8:**
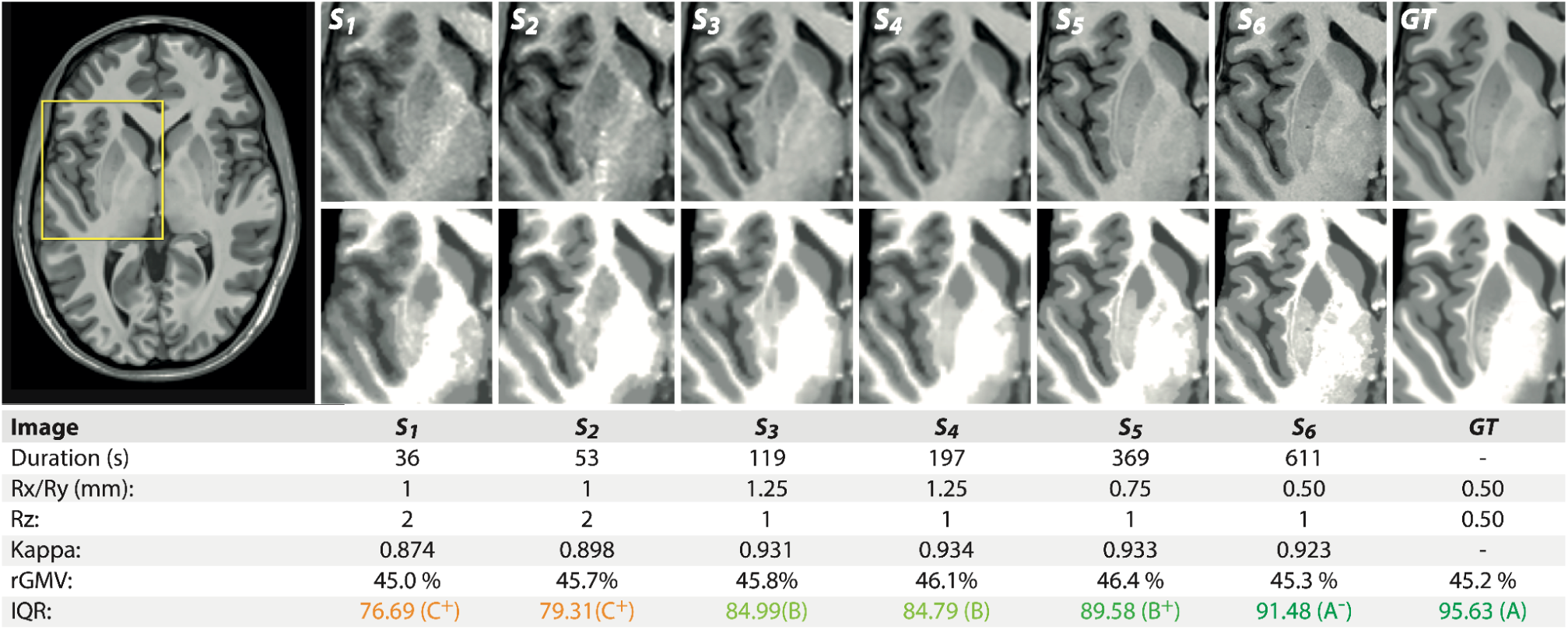
The test-retest sample analysis focuses on 6 images with an increasing scan-time and image quality. The top and bottom rows represent the intensity normalized T1 images and the CAT12 segmentation, respectively, with images/segmentations progressively converging (higher kappa) to the average gold-standard (created from the best 127 test-retest images). The relative GM volume (rGMV) is slightly underestimated in low quality data.

## Discussion

Here we introduced a QC framework for structural (T1-weighted) MRI data. We defined and validated various automated quality ratings based on well-defined BWP image quality features such as noise, inhomogeneity, and resolution (Aubert-Broche et al., 2006) and integrated them into a single useful SIQR score to facilitate practical applications in context of clinical and cognitive neuroscience. We further demonstrated that our measures (i) are robust to simulated segmentation problems and cortical atrophy; (ii) are independent from sex, and brain size, showing only minor expected associations with chronological age, as well as severe disease-related changes; and (iii) allow the reliable assessment of motion artifacts within a protocol. In artifact-free data, image quality typically varies between 2.5 and 5 rps (0.25-0.5 grade), whereas light or strong artifacts typically result in a reduction of ratings by 5 or 10 rps (equivalent to a 0.5 or 1.0 increase in grades), respectively. T1-weighted images with low quality ratings might show a systematic underestimation of gray matter volume, first demonstrated in case of motion artifacts by Reuter et al. (2015). Strongly affected data should therefore be excluded from the analyses. In case of less severe artifacts, the quality rating might be included as a covariate (Garcia-Dias et al., 2020; Pomponio et al., 2019) or using weighted least squares (Lutti et al., 2022) during statistical analyses. However, an empirical comparison of the complex statistical effects of alternative approaches to account for automatically generated (i.e. known) quality differences in downstream analysis tasks is still lacking.

The proposed QC framework offers a simple and efficient approach to identify structural MRI scans that are suitable for the prospective use in structural processing tools and brain imaging analysis in both clinical and research settings. This was also confirmed in previous studies by Gilmore et al. (2021), Hoffstaedter et al. (2024) and Ma et al. (2022), that evaluated the utility of earlier versions of this QC framework.

In the following sections, we discuss further aspects of the development of our SIQR measure and its subordinate quality measures (with regard to existing alternatives), their performance in simulated and real data samples, and their potential for assessing the quality of images from other sequences and modalities.

### SIQR measure development

Image quality measures are commonly estimated from the image background (Mortamed et al., 2008; Esteban et al., 2017). In contrast, our approach focuses on estimating quality only within the brain for three reasons. First, the background values in public datasets can be corrupted by various defacing and skull-stripping routines (Bhalerao et al., 2022, Rubbert et al., 2022). Second, the background may contain artefacts (e.g., motion artefacts from the jaw or tongue) or unwanted properties (e.g., noisy backgrounds in MP2RAGE; Marques et al., 2010) that do not necessarily affect the brain, or conversely, the artefacts in the brain do not/are less likely to affect the background. Third, the background does not provide information about image inhomogeneity, tissue contrast and spatial anatomical resolution (Likar et al., 2001). Furthermore, the evaluation of image quality within tissues must take into account structural aspects such as (i) the partial volume effect, where a voxel contains tissue of more than one tissue class, and (ii) changes in brain development and aging, such as tissue degeneration due to white matter hyperintensities, small vessel disease or perivascular spaces (Westlye et al., 2010; Lynch et al., 2024). Consequently, the proposed framework adapts these regions of interest by applying specific thresholds and morphological operations to minimize bias from age/disease, as we have demonstrated in IXI and ATLAS datasets. Moreover, the proposed intensity-based measures are normalised by (minimum) tissue contrast rather than signal intensity, as this is more relevant for segmentation and surface reconstruction (Ashburner et al., 2005).

Our proposed individual quality subscores have largely been established based on well-known image quality aspects of the BWP, which was built to represent the large variability in image quality of structural T1-weighted MR images (Aubert-Broche et al., 2006; Luo et al., 2022; Tönnes et al., 2024). By taking into account these predominant aspects of image quality, we have created ratings that are easy to understand, even without a technical background. The ratings were integrated into a single SIQR rating to support the users during the evaluation process. To combine the measures, we have used an RMS-weighted average (of the grades) with a power of 4 rather than 2, to place greater emphasis on the more problematic aspects of image quality. This is relevant because effects of severe problems can often not be compensated by other factors, e.g. if there are severe motion artefacts, a much higher image resolution can typically not account for this.

In particular, SIQR is strongly predictive of segmentation accuracy (quantified by the kappa measure) and the extracted GM volume although its quantification is largely independent from structural features. Thus, SIQR can facilitate the estimation of image quality-related variance in individual scans or samples even for non-expert. Alternative quality control tools, such as MRIQC (Esteban et al., 2017), might be challenging for novices due to non-standardized measures that require substantial user experience. Moreover, a normalisation using BWP quality features also enables a direct comparison across protocols (see test-retest example above), though caution is advised, as the results may be subject to bias by (i) our focus on a segmentation-centred definition of quality, (ii) the population under study and (iii) project-specific needs or considerations (e.g., optimised MR parameters to image specific structural changes rather than pre-processing).

### Identification of scans with data anomalies and artifacts

The proposed framework is part of the CAT12 preprocessing and utilizes the CAT12 segmentation, but could also be used as an independent SPM batch with other segmentation algorithms, e.g. from SPM (Ashburner et al., 2005) or SynthSeg (Billot et al., 2023). Segmentation routines are widely used for structural brain analyses and have undergone intensive testing to be valid, accurate and robust for a variety of protocols, individual anatomies, and demographics (e.g., Ashburner et al., 2005, Mendrik et al., 2015, Gaser et al., 2024), making them ideal for image quality analysis. By focusing on general global aspects of the scan rather than local ones, problematic structures and areas such as partial volume effect voxels or WM lesions can be omitted, allowing precise, robust, and largely consistent results even in case of severe classification faults (e.g., failed skull-stripping or miss-classification), as tested here under simulated conditions.

Although SIQR could be used for fully automatic outlier detection (see also Gilmore et al. (2021) and Bhalerao et al. (2024)), we believe that the huge variability of type of artefacts, their regional occurrence and their impact on image processing still requires study-specific knowledge and, if possible, a short user inspection. For instance, in cases when locally limited or mild artefacts affect regions that are not relevant to the study (e.g., if the study focuses on frontal regions, cerebellar artefacts from jaw movements are acceptable) or whenever lower preprocessing accuracy is acceptable (e.g., for local alignment of brain surfaces or atlases for other modalities).

Multivariate outlier detection schemes that are typically applied based on the processed data of a sample in the normalized feature space, using similarity analysis of normalized GM data (e.g. the Gram matrix or kernels) in CAT12 (see section <u>Software</u>), can be used to detect outliers with preprocessing problems or highly deviating anatomy. However, the proposed image quality assessments are specifically designed to measure differences of image quality (in native space) rather than segmentation accuracy (in normalized space) or anatomical properties, such as stroke lesions, and can therefore be used in addition to previously mentioned outlier detection schemes to identify cases where image artifacts could bias analysis.

### The role of subordinate quality ratings

While the SIQR composite is sufficient for most analyses, our framework facilitates deeper insights providing more specific subordinate ratings.

The *noise-to-contrast ratio* (**NCR**) is not only a very robust measure as it can be quantified in different regions, it is also very sensitive to motion artefacts and gives the most relevant values when the image resolution is adequate (<1.5 mm). In our comparisons NCR supports a slightly better artifact separation than the best performing MRIQC measures, such as *snr_wm*, *snr_total, cjv* and *cnr,* which also evaluate noise differences.

The *inhomogeneity-to-contrast ratio* (**ICR**) had little to no impact on detecting problematic data (e.g., when testing for segmentation quality kappa in the BWP or motion artifacts in MR-ART), as inhomogeneities tend to describe more protocol/scanner specific aspects and can be corrected fairly well in most protocols (Belaroussi et al., 2006). Increased inhomogeneities typically occur in high-field scans without protocol specific correction schemes. Although possible disadvantages are generally outweighed by superior resolution and higher signal-to-noise ratio, bias correction schemes in preprocessing routines can fail in some cases (Feinberg et al., 2023). It is therefore recommended to retain this measure in a general rating.

The *resolution score* (**RES**) rates the voxel size in terms of how good structural features can be imaged. Nevertheless, structures can still be biased by interpolation (Tian et al. 2021), blurring, noise or motion artifacts. A real quantification of the sharpness of anatomical structures by our *edge-to-contrast ratio* (**ECR**) is therefore essential, although it is strongly affected by noise and the segmentation quality compared to other ratings.

*The full-brain Euler characteristic* (**FEC**) represents our adaptation of surface topology (Backhausen et al. 2016, Rosen et al., 2018). The measure showed strong association with noise levels and supports the identification of motion artifacts. However, compared to NCR it is more noisy and depends strongly on the input segmentation and MR protocol. Data with low spatial resolution or faulty/simplified segmentations with limited amount of details can have less defects and result in better ratings. Nevertheless, FEC presents a good extension to the NCR and ECR measurements.

In contrast to MRIQC (Esteban et al., 2017) that provides a variety of raw unscaled measures (with reversely signed scored ones among them), we tried to establish measures that reflect the known specific perturbances and are directly interpretable by applied scientists. All QC measures can be used in statistical analyses or machine learning models according to the study needs (Bhalerao et al., 2024).

### Evaluation in simulated and real data

Simulated data allow basic validation of methods under expected conditions and comparison with actual ground truth results. The BWP is a standard for evaluation of structural brain image preprocessing (Luo et al., 2022; Tönnes et al., 2024) and was used here to define and normalise our quality measures and to test their relationship with the segmentation accuracy. Due to the robustness of CAT12, we decided to simulate extreme segmentation problems. The results showed high stability for the NCR and ICR measures, but high susceptibility to error for the ECR and FEC in the case of severe over/underestimation of tissue segments, as both depend on the correct definition of the GM/WM boundary. In the simulated aging phantom (Rusak et al., 2021), the results showed small systematic but negligible changes in the quality measures, which could also be due to small differences in the simulated images (see bias differences in Figure 5F). Since a lot of our tests relied on BWP, which is limited in its ability to simulate artifacts or new protocols, such as MP2Rage, new frameworks such as TorchIO (Pérez-García et al., 2021) present a possible next step for future tests. Nevertheless, an empirical validation on real MRI datasets was necessary to demonstrate the validity and practical benefits of the introduced quality assessments and to avoid over-adaptation to synthetic data. Therefore, we used the IXI and ATLAS datasets to demonstrate that SIQR is unaffected by age, sex, head size, or severe structural disease-related changes. In MR-ART, we tested the ability to identify different degrees of motion artifacts and the effects on gray matter segmentation in aging. Overall, we demonstrated the robustness and applicability of our SIQR measure.

### Shortcomings and outlook

The QC measures proposed in this study were designed to be independent from segmentation accuracy and were tested in a variety of protocols. However, useful results can only be expected for valid segmentation inputs (where we focus on CAT12), and highly specific T1-weighted protocols may result in unexpected ratings. Other modalities such as T2-weighted, proton density weighted or FLAIR images can be assessed, however, the dependance of the SIQR on separability of CSF, GM and WM may result in low quality scores. In addition, our ratings are not designed to assess functional or diffusion data where more specific tools are available (Christodoulou et al., 2013; Roalf et al., 2016; Nakua et al., 2023). Although such data can also be used for tissue segmentation, the low GM-WM contrast is challenging and the resulting segmentations or surfaces are less accurate and possibly biased compared to typical T1-weighted images (Ashburner et al., 2005). It is important to note that preprocessing tools are designed to work reliably even on problematic datasets, and that results from these images can often still be used, though these should be interpreted with more caution. Moreover, scanner-specific changes, such as geometric distortion, have not been considered. Consequently, our measures are not designed to monitor scanner properties that require real MRI phantoms (Belli et al., 2016; Davids et al., 2014).

In addition, our scan-rescan results demonstrated instances of comparable segmentation quality with up to 40% faster scan-times. This is particularly relevant for clinical MRI, where cost-effectiveness (short scan times for high patient throughput) presents an essential aspect, and images of adequate, but not exceptional, quality are appropriate for diagnosis (Jhaveri et al., 2015; Rofsky et al., 2015), simultaneously reducing both financial and environmental costs (Chaban et al., 2024). On the other hand, using only adequate image quality for certain projects does not eliminate the need for cutting-edge resolution (Feinberg et al., 2023), though improperly enhanced image resolution, e.g., 0.5×0.5×1.5 mm for 1.5 Tesla systems, often leads to increased noise or parallel imaging artifacts that can disturb preprocessing. It is therefore advisable to pilot modified protocols for the preprocessing pipelines you plan to use or follow the established standard protocols, e.g., ADNI (Arani et al., 2024) and HCP (Van Essen et al., 2013).

## Conclusion

Our fully automatic quality control framework enables a standardized, accurate, and robust evaluation of large heterogeneous datasets to detect outliers with inadequate image quality using a single image quality rating SIQR. Its flexibility, low-cost, and simplicity support a wide range of applications and can provide a valuable contribution to quality assurance in clinical practice and research.

## List of abbreviations

BWP: Brain Web Phantom
CAT: Computational Anatomy Toolbox
CSF: CerebroSpinal Fluid
GM: Gray Matter
ICR: Inhomogeneity Contrast Ratio
NCR: Noise Contrast Ratio
MRI: Magnetic Resonance Imaging
SPM: Statistical Parametric Mapping
RES: Root mean square Resolution Rating
ECR: Edge Contrast Ratio
FEC: Full-brain Euler Characteristic
QC: Quality Control
SIQR: Structural Image Quality Rating
TIV: Total Intracranial Volume
WM: White Matter

## Acknowledgement

We want to thank Dr. Daniel Gülmar for his helpful comments, Benjamin Thyreau for the scan-rescan dataset, and all other projects providing publicly available data (even if not presented here). This manuscript reflects the views of the authors and may not reflect the opinions or views of the different projects.

IXI was made possible by the research grants of action charity, the Engineering and Physical Sciences Research Council (EPSRC GR/S21533/02), and the Medical Research Council.

This work was funded by the *Deutsche Forschungsgemeinschaft* (DFG) Nr. <u>417649423</u> and grant 351849 from the Research Council of Finland under the frame of ERA PerMed (“<u>Pattern-Cog</u>”).

## Data and materials

MRI data is available by the original providers given in Table 1. The QC functions are part of the CAT12 toolbox (Gaser et al., 2024, https://neuro-jena.github.io/cat/).

## Availability and requirements

- Project name: (Quality Metric of the) Computation Anatomy Toolbox (CAT)
- Project home page: https://neuro-jena.github.io/cat/
- Operating system(s): Linux, Mac OS, Windows
- Programming language: MATLAB™
- Other requirements: Statistical Parametric Mapping (SPM), R7771 https://www.fil.ion.ucl.ac.uk/spm/
- License: GNU GPL version 2 or higher
- RRID: SCR_019184
- Bio.tools ID: -

Processing was done under Mac OS using MATLAB 2023a, SPM12 R7771, CAT12.8.2 R2166 (segmentation) and CAT12.9 R2647 (quality control).

## Data and Code Availability

Data used in the preparation of this manuscript are available publicly (might require registration and approval). The framework is part of the CAT12 toolbox, the code used to generate the results and figures is available on request from the first author.

## Ethics Statement

The study relies on publicly available datasets that were collected by complying to ethical standards.

## Authorship Contributions

**Robert Dahnke:** Methodology, Software, Validation, Formal analysis, Investigation, Resources, Data Curation, Writing - Original Draft, Writing - Review & Editing, Visualization, Project administration. **Polona Kalc:** Writing - Review & Editing. **Gabriel Ziegler:** Writing - Review & Editing. **Julian Großkreutz:** Writing - Review & Editing, Funding acquisition. **Christian Gaser:** Methodology, Software, Writing - Review & Editing, Supervision, Project administration, Funding acquisition.

## Supplement

### Technical Definition of Quality Measures

#### Scaling

Simple linear scaling function:

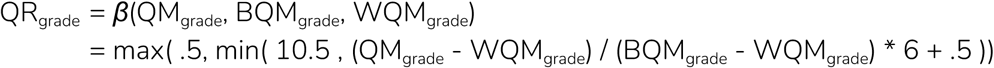

#### Noise

The first quality rating characterizes image noise defined here as *noise-to-contrast ratio* (**NCR**), to describe how well tissue can be locally separated independent of the protocol-specific tissue contrast. The estimation was specified as the minimum of the average local standard deviation, Õ, of the bias corrected image C_bc_ within the optimized *white matter* (**WM**) and *cerebrospinal fluid* (**CSF**) regions WMe and CSFe. The values were normalized by the minimum tissue contrast c_min_ and scaled by the results of the linear fit:

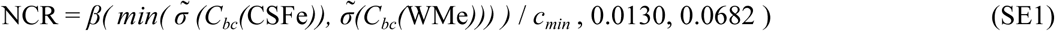

with 0.0130 as the best, and 0.0682 as the worst rating of the unscaled measure obtained for the BWP train dataset. For data analysis, the bias corrected image C_bc_ allows for a more meaningful characterization of the local varying noise level than the original image, since it considers processing problems in areas with low signal intensity and increased noise. The local standard deviation Õ was estimated in a 5×5×5 voxel neighborhood of a voxel and averaged to reduce the influence of remaining inhomogeneities.

The CSF and WM regions, rather than the background, were used because the background can contain interferences that do not affect the brain (Kruggel et al., 2010; Marques et al., 2010), or could be affected by anonymization of subject-features by defacing or brain extraction (Figure 2A). CSF and WM are beneficial for noise estimation compared to the *gray matter* (**GM**) because (i) they cover relatively large and homogenous areas and (ii) are less affected by partial volume effects and locally varying tissue contrast (e.g. by myelination). However, using only CSF regions often failed in younger subjects and low-resolution data, while the exclusive use of WM led to age-related effects caused by WM lesions or small vessel disease or perivascular spaces. The regions are optimized by an erosion step and additional tissue thresholds to avoid side effects by partial volumes, segmentation method, or WM lesions in elderly subjects that are quite similar to noise or artifacts (Figure 2B). The minimum tissue contrast c_min_ between CSF, GM, and WM was used because a greater GM-WM contrast led to problems in detecting the CSF-GM and CSF-background boundaries.

#### Inhomogeneity

In order to assess intensity inhomogeneity in images (often referred to as bias), the *coefficient of joint variation* (**CJV**; Likar et al., 2001) proved to be one of the most suitable measures (Belaroussi et al., 2006):

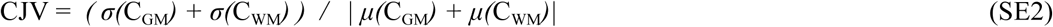

However, since it is known that the GM is strongly influenced by partial volumes and locally different GM intensities (Westlye et al., 2010), only the standard deviation σ of the WM is determined here. Similar to the NCR, the minimal tissue contrast is used rather than the GM-WM contrast. To remove noise driven variance, a Laplacian filter with Dirichlet boundary condition is applied in the WMe area, resulting in a locally averaged image Cs, which was used to estimate the *inhomogeneity-to-contrast ratio* (**ICR**):

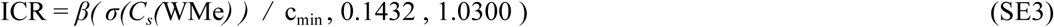

Since most methods are able to correct strong inhomogeneities almost without loss of segmentation accuracy (e.g., approach X in Figure 1B; Belaroussi et al., 2006), a weaker weighting was used. The worst BWP inhomogeneity level describes a grade C (see Figure 3) that can be already measured in 3 Tesla data without protocol-based corrections.

#### Resolution

The spatial resolution of MRI images plays an important role in obtaining meaningful representations of anatomical structures. For the general assessment of voxel volume and proportion in a single value, we use the RMS notation to define the *RMS voxel resolution* (**RES**):

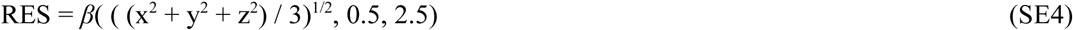

As a consequence of this definition, outliers with exceptionally low resolution in one of the three dimensions are weighted much higher than outliers with high resolution, resulting in an asymmetric evaluation where similar (isotropic) resolutions are preferred. The quality range was arbitrarily determined to characterize typical resolutions, with a simple scaling step size of 1 grade (10 rps) for another 0.5 mm, with 0.5 mm as an excellent result and 2.5 mm as the lowest quality limit close to the average cortical thickness in humans.

For the principal evaluation, we tested RES by reducing and re-interpolating the tissue label map of the BWP to quantify the loss of information by Cohen’s kappa (Cohen, 1960). RES yielded higher Spearman correlation coefficient than the simple voxel mean RESM (rho_RES_ = 0.994; rho_RESM_ = 0.965; with RESM = *β*((x + y + z)/3, 0.5, 2.5). Although RES provides a good description of resolution under normal conditions, it has the major limitation that it does not quantify the true anatomical level of detail, i.e. how well fine structures are defined and how sharp the boundaries are.

The *edge-to-contrast ratio* (**ECR**) is the average gradient Δ of the GM/WM boundary (outlined by the segmentation and masked for extreme gradients, e.g. between CSF and WM and blood vessels) and normalized by the minimum tissue contrast and scaled similarly to the RES rating. It allows an evaluation of structural resolution independent of resampling or smoothing.

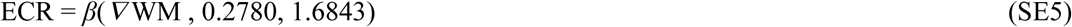

To test the quantification of anatomical rather than image resolution, spatial details were removed by resampling (downsampling to 1.25:0.25:3.00 mm and resampling to 1.00 mm) and smoothing (0:0.25:3.00 mm) a BWP image with 1% noise, 20% inhomogeneity of field A, and 1 mm resolution. The parameter test range was defined by the resolution of the BWP, the minimum smoothing resolution (0.2 mm for 1.0 mm data) and the average cortical thickness of 2.5 mm. The resulting images were then segmented to quantify changes using Cohen’s kappa. In both cases, the final voxel resolution remains constant, so that the voxel-based RMS resolution measurements are identical even though the images become blurred and kappa decreases (see Figure S1). In contrast, our new ECR measure allows quantification of both test cases, although quantification of GM/WM edge strength and tissue contrast introduces further variance (r>0.98, p<7e-07).

However, there are several limitations of the measure itself, but also of the test design: (i) the BWP is limited in its anatomical details, supporting only 1 mm resolution with some partial volume effect, (ii) linear/spline resampling and smoothing affect the measures differently, (iii) kappa only quantifies segmentation accuracy, but not the quality of more complex surface reconstruction (e.g. Hausdorff distance to the GT surface) that could be used. Nevertheless, ECR already represents a significant step forward in quantifying image detail in real data.

**Figure S1:**
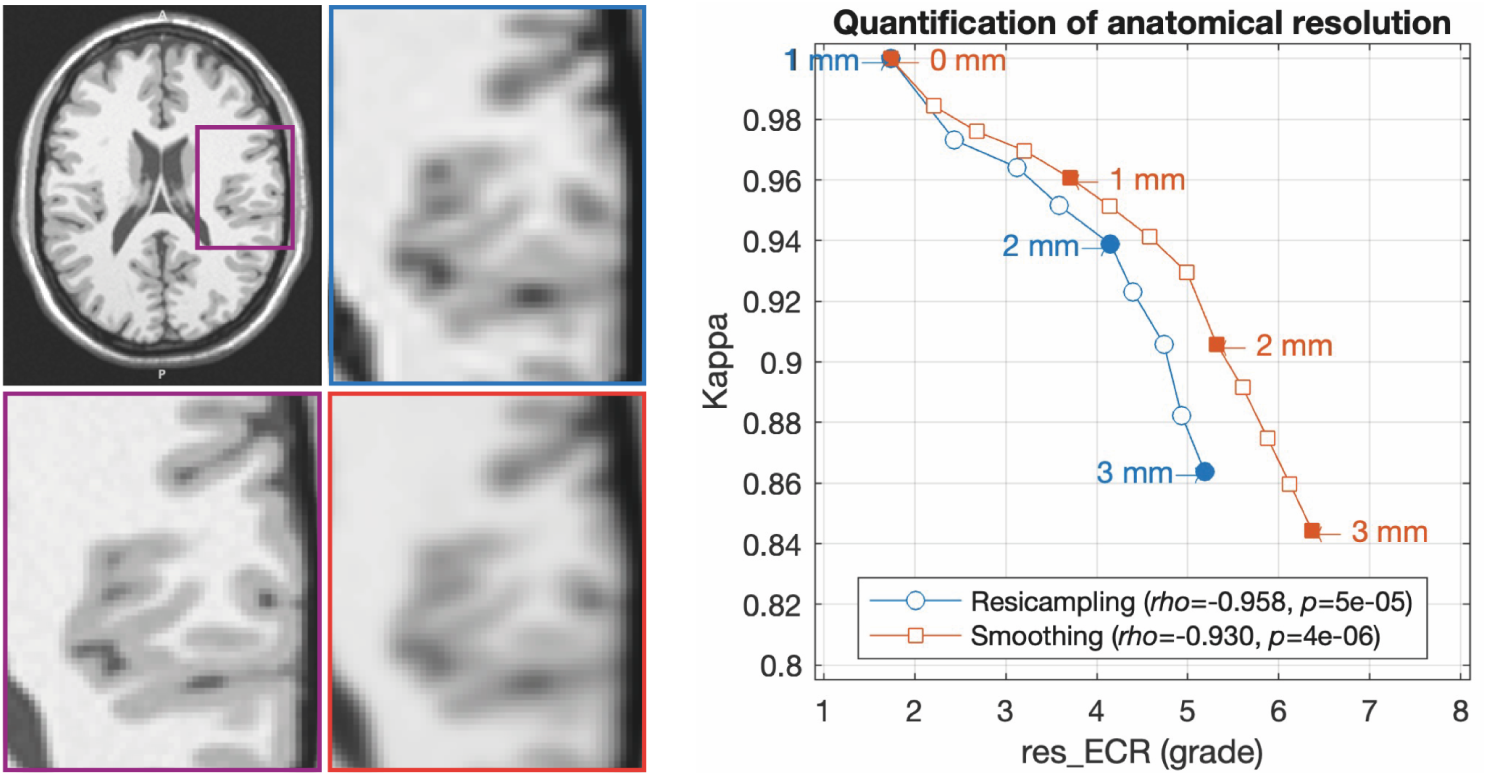
Shown are the effects of the simulated reduced resolution by resampling (downsampling to low resolution and followed upsampling to original resolution) and Gaussian smoothing on the segmentation accuracy (quantified by kappa) for the *edge-to-contrast ratio* (**ECR**) resolution measure. Quantifying only by voxel resolution (RMS resolution score) would give the same value (grade 2 for 1 mm data) for all test cases (results for the BWP with 1% noise, 20% inhomogeneity field A and isotropic 1 mm resolution). It is also obvious that there is a large step between the original resolution and the first resampled resolution, which describes the general information loss of data resampling and would be different in real data or if input data with higher ground truth resolution were used (i.e. test bias). In addition, quantification in low resolution data (below 2 mm) becomes increasingly difficult due to the highly folded and thin cortical band (sampling theorem).

#### Surface topology

In order to approximate the surface topology in a reasonable time, the *full-brain Euler characteristic* (**FEC**) was estimated at a resolution of 2 mm. The whole-brain WM surface was used rather than the typical neocortical hemispheres of most surface pipelines. To account for partial volume effects at the lower resolution, two WM surfaces were generated at thresholds of 0.25 and 0.75. As we observed more defects in children due to the thin developing WM structures, we used a maximum filter to extend and stabilize the surface creation.

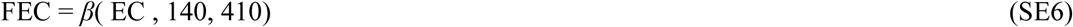

where the Euler characteristic EC is defined as EC = V - E + F with V as number of vertices, E as the number of edges, and F as the number of faces of the created brain surface.

### Detailed BWP Results

As CAT produced highly accurate segmentations even in problematic BWP cases, we simulated severe segmentation errors to test the robustness of our measures in exceptional situations. We simulated problems in skull stripping and tissue over/underestimation in 15 BWP cases with 1 to 9% noise, 1 mm resolution, and 20% inhomogeneity of 3 fields. To test the influence of preprocessing problems, we simulated severe distortions due to skull-stripping (i.e., missing brain/additional head) and tissue segmentation errors (e.g., WM overestimation/underestimation). To simulate inaccurate skull-stripping, a strongly smoothed map of random values and distance information was used to add/remove tissues to the segmentation. We then modified the tissue boundaries by eroding/dilating the tissue segments, using a maximum/minimum filter to preserve the partial volume effect (dWM, eWM, eCSF, dCSF, eWM & eCSF). Although these changes do not affect the kappa values as much as skull stripping, the changes in GM volume are extreme and comparable to thickness changes of approximately 1 mm (Figure S2B). In all cases, the variance of the measures was positively correlated with the noise of the BWP, i.e. quantification of noisy images is again more error-prone. Nevertheless, most of our measures are quite robust even to severe errors with kappa values below 0.5. Finally, this also means that our image quality measures do not (directly) quantify the quality of the preprocessing itself (i.e., segmentation accuracy). However, to the extent that lower image quality leads to lower processing quality, it can be assumed that processing accuracy is generally lower for low quality input and that further user intervention is required (e.g., to check and remove problematic scans).

**Figure S2:**
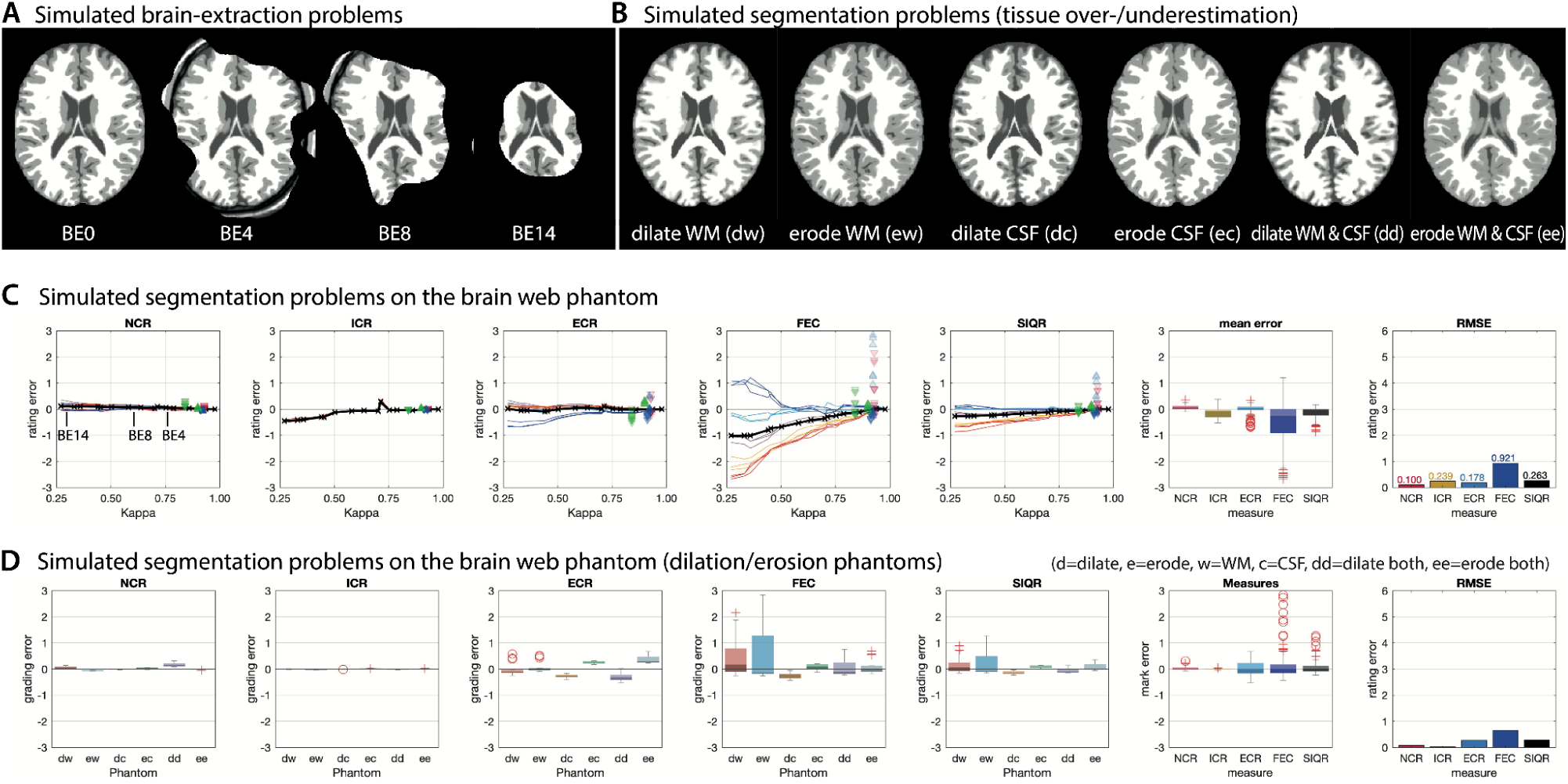
We simulated the effects of segmentation accuracy for brain extraction problems (A) and for tissue over- and underestimation (B). Effects of skull-stripping problems are shown as lines in C, while the segmentation problems are shown as scatter. Our quality scores are almost unaffected by severe skull stripping problems (kappa < 0.6), as even small regions are sufficient for global estimation. Tissue classification errors are more challenging even if the reduced accuracy is quite small because the contrast estimation is biased. In general, problematic cases tend to underestimate the image quality, which indirectly helps to identify serious pre-processing problems.

### Detailed Aging Phantom Results

**Figure S3:**
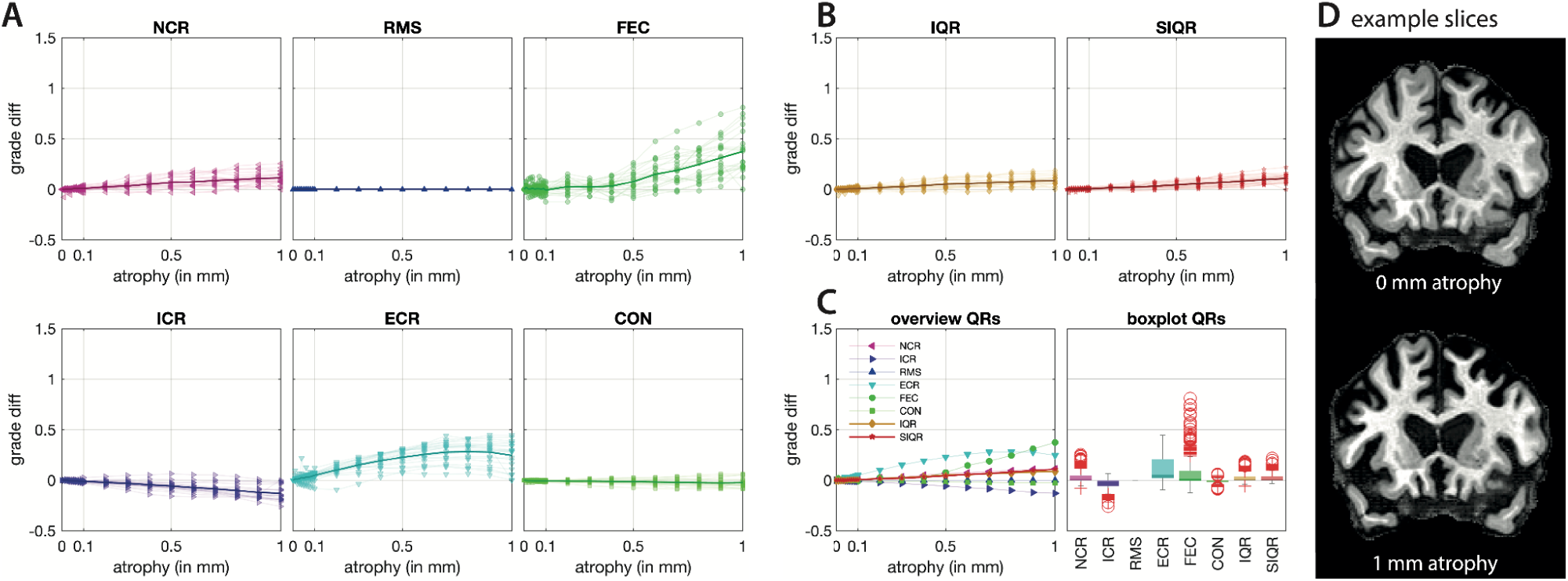
Changes in current quality score (A) and combined measures (B) for the phantoms presented in (Rusel et al., 2021, D), which simulated neocortical atrophy of up to 1 mm in 20 subjects from the ADNI database, where 0.01 mm cortical loss is equivalent to 1 year of healthy ageing. As only the neocortical thickness is changed (D), similar measurements are expected regardless of the simulated atrophy rate. However, the slightly different inhomogeneity and noise pattern in the CSF could introduce further bias in the evaluation.

### Detailed Real Data Results - IXI

**Figure S4:**
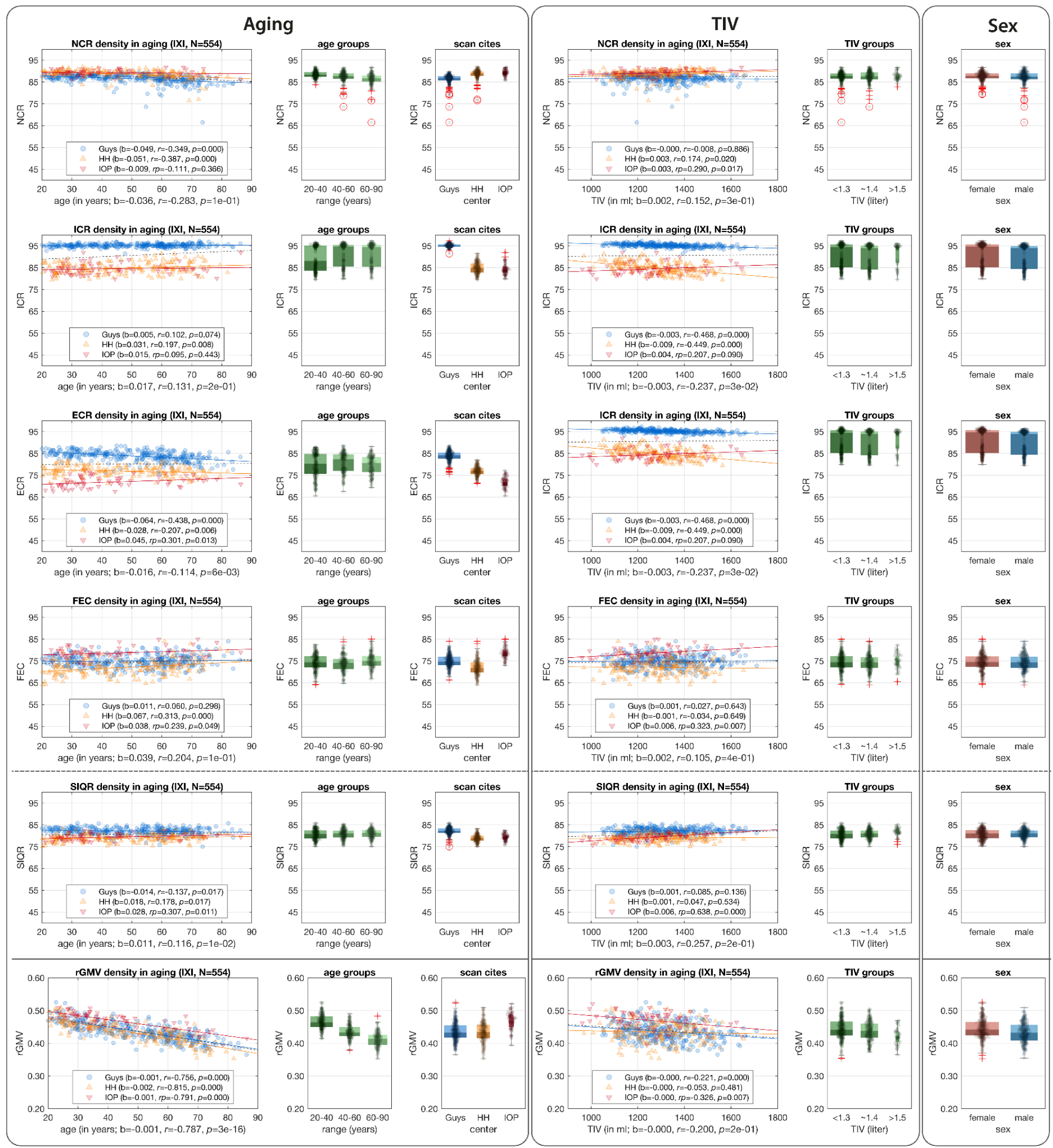
Shown are the changes of all quality ratings (NCR, ICR, ECR, FEC, and SIQR) and the relative GM volume (rGMV) for aging (left), total intracranial volume (TIV, center), and sex (right).

### Detailed Real Data Results - MR-ART

**Figure S5:**
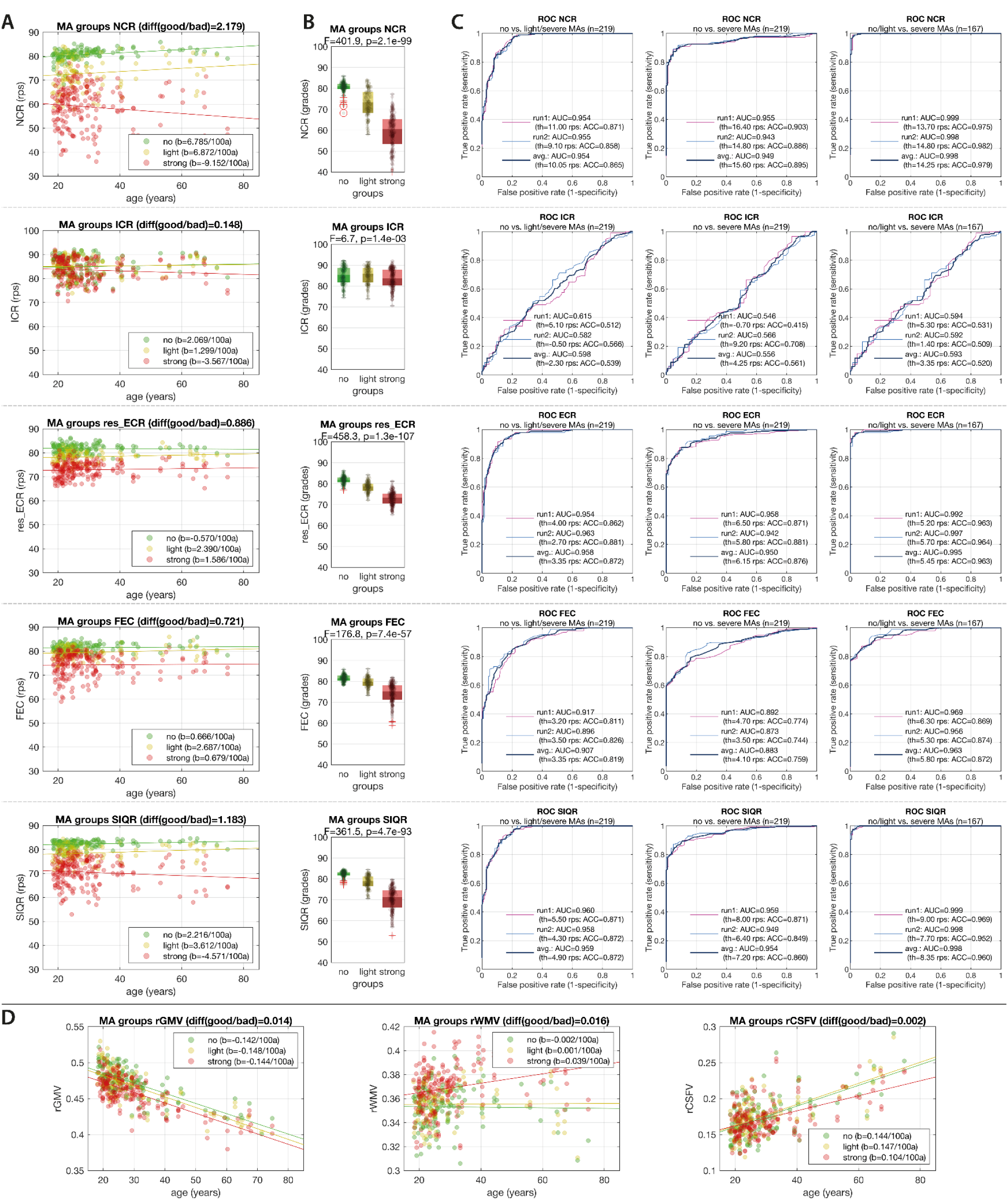
Shown are the changes of all quality ratings (NCR, ICR, ECR, FEC, and SIQR) in aging (A) and grouped by the expert rating (B). (C) shows the ROC analysis of each quality rating to separate the expert rated motion groups. The change of the *relative GM volume* (**rGMV**) for aging is shown in (D).

**Figure S6:**
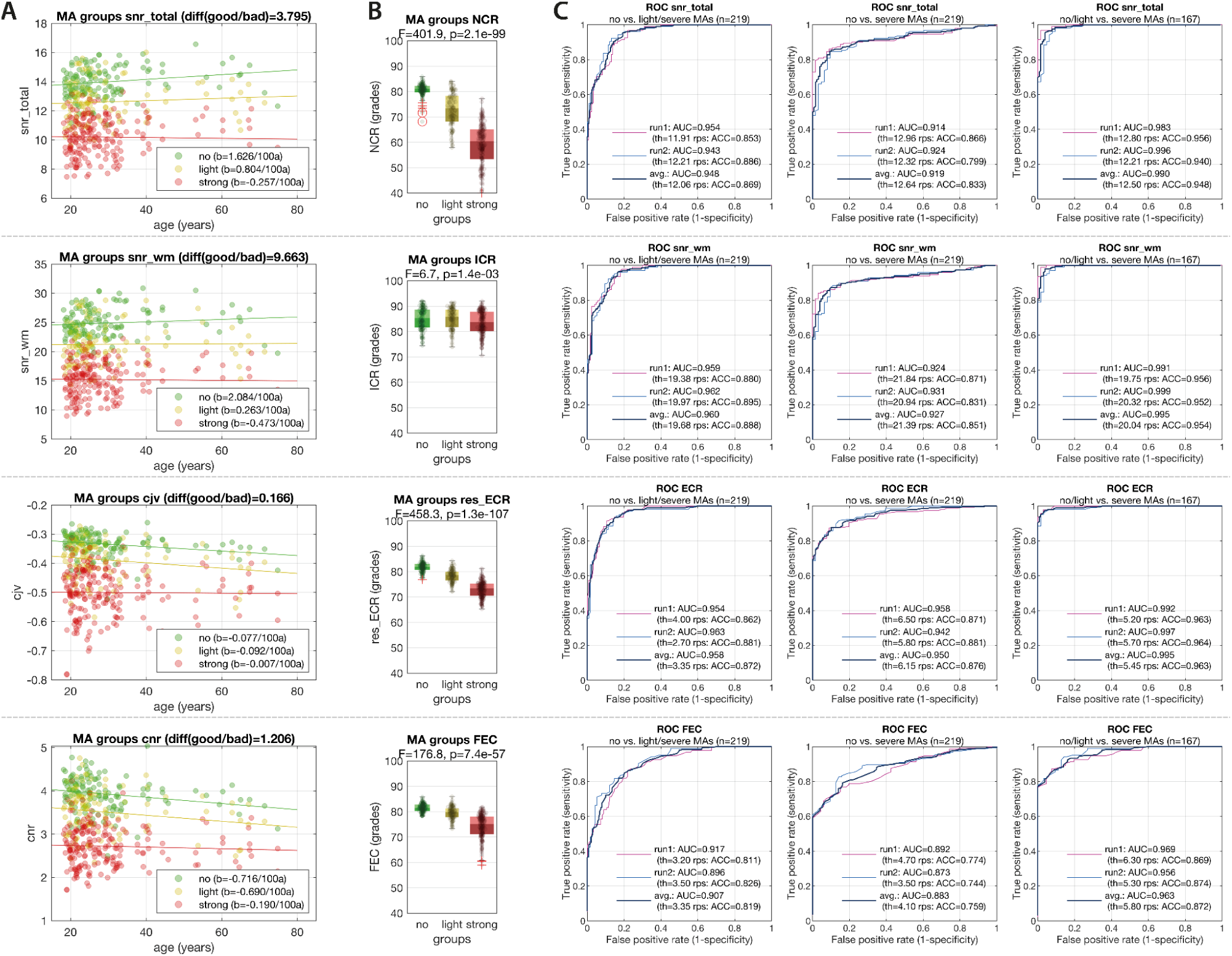
Shown are the changes of the four best performing MRIQC quality metrics (snr_total, snr_wm, cjv, cnr) in aging (A) and grouped by the expert rating (B). (C) shows the ROC analysis of each quality rating to separate the expert rated motion groups.

